# Microglia inhibition rescues developmental hypofrontality in a mouse model of mental illness

**DOI:** 10.1101/254656

**Authors:** Mattia Chini, Christoph Lindemann, Jastyn A. Pöpplau, Xiaxia Xu, Joachim Ahlbeck, Sebastian H. Bitzenhofer, Christoph Mulert, Ileana L. Hanganu-Opatz

## Abstract

Cognitive deficits, core features of mental illness, largely result from dysfunction of prefrontal-hippocampal networks. This dysfunction emerges already during early development, before a detectable behavioral readout, yet the cellular elements controlling the abnormal maturation are still unknown. Combining *in vivo* electrophysiology and optogenetics with neuroanatomy and pharmacology in neonatal mice mimicking the dual genetic - environmental etiology of psychiatric disorders, we identified pyramidal neurons in layer II/III of the prefrontal cortex as key elements causing disorganized oscillatory entrainment of local circuits in beta-gamma frequencies. Their abnormal firing rate and timing result from sparser dendritic arborization and lower spine density. Pharmacological modulation of aberrantly hyper-mature microglia rescues morphological, synaptic and functional neuronal deficits and restores the early circuit function. Elucidation of the cellular substrate of developmental miswiring related to later cognitive deficits opens new perspectives for identification of neurobiological targets, amenable to therapies.

**Highlights:**

- Mice mimicking the etiology of mental illness have dysregulated prefrontal network
- Structural and synaptic deficits cause abnormal rate and timing of pyramidal firing
- Weaker activation of prefrontal circuits results from deficits of pyramidal neurons
- Rescue of microglial function restores developing prefrontal circuits

## INTRODUCTION

Cortical function relies on the precise wiring and activation of diverse populations of pyramidal cells and interneurons that are entrained in oscillatory rhythms. While recent studies have revealed several assembling rules of cortical microcircuits in the adult brain^1,2^, their ontogeny is still poorly understood. Given the uniqueness of developing brain in its spatial and temporal organization of coordinated activity^3–5^, the depolarizing action of GABA^6,7^ and the formation of transient connectivity patterns^8,9^, the functional coupling within immature microcircuits is likely to bear equally unique traits. Elucidating the features of such immature networks is of paramount importance in the context of neurodevelopmental disorders^10^, as their early disruption is thought to underlie the later emersion of the devastating symptoms that characterize these diseases^11^.

We have recently begun to elucidate the mechanisms of functional coupling within developing brain and shown that layer II/III pyramidal neurons in the prefrontal cortex (PFC) of the neonatal mouse play a fundamental role in generating beta/low gamma oscillations^12^. At adulthood, coordinated activity in gamma frequency band is instrumental to cognitive processing^13,14^ and relates to the pathophysiology of psychiatric disorders^15–18^. Disturbed gamma activity has been observed long before the onset of psychosis in high risk humans^19^ and during neonatal development in animal models^20^. However, the circuit dysfunction underlying such abnormalities is still fully unknown.

To address this knowledge gap, we interrogate the developing prefrontal network in a mouse model mimicking the genetic (mutation of Disrupted-In-Schizophrenia 1 (DISC1) gene) and environmental (challenge by maternal immune activation (MIA)) etiology of mental disease (dual-hit GE mice). At adult age, these mice reproduce to a large extent the behavioral abnormalities identified in human psychiatric disorders, such as elevated anxiety, depression-like responses, memory and attention deficits as well as abnormal social behavior^21,22^. To elucidate the mechanisms of developmental dysfunction we focused on neonatal age (end of 1^st^ – beginning of 2^nd^ postnatal week) of rodents that roughly corresponds to the second trimester of human pregnancy, a period of high vulnerability for mental disorders^23^. We combine *in vivo* and *in vitro* electrophysiology, optogenetics, and pharmacology with confocal microscopy-based structural investigations of the prelimbic cortex (PL), the prefrontal sub-division strongest coupled with the hippocampus^24,25^. We show that pyramidal neurons in layer II/III exhibit major morphological, synaptic, and functional deficits and lack the ability to organize the beta-gamma entrainment of local prelimbic circuits in neonatal dual-hit GE mice, while layer V/VI neurons are largely unaffected. Pharmacological intervention restoring the aberrant hyper-maturation of microglia in dual-hit GE mice rescues layer-specific electrophysiological and structural deficits of developing prefrontal networks. These data elucidate the mechanisms of developmental miswiring related to mental illness and highlight the tight link between neuronal and glial dysfunction.

## RESULTS

### Layer- and frequency-specific dysfunction of local circuits in the prelimbic cortex of dual-hit GE mice

To get first insights into the source of prelimbic dysfunction in dual-hit GE mice, we performed extracellular recordings of the local field potential (LFP) and multiple unit activity (MUA) over prelimbic layers using four-shank 16 site-electrodes in postnatal day (P) 8-10 control (n=38 pups from 13 litters) and GE mice (n=18 pups from 6 litters). This developmental stage corresponds to the initiation of hippocampus-driven entrainment of prelimbic circuitry^5,12^. The exact position of recording sites covering layer II/III and layer V/VI was confirmed by the reconstruction of electrode tracks *post-mortem* (Fig. 1a). In line with our previous findings^5,20,26,27^, the first patterns of network activity of the neonatal PL in all investigated control and dual-hit GE mice were discontinuous, i.e. spindle-shaped oscillations switching between theta and beta-gamma frequency components alternated with long periods of network silence (Fig. 1b). The firing of prelimbic neurons is strongly timed by the oscillatory rhythms. As previously reported^27^, the patterns of network oscillations and neuronal firing in the PL were similar in urethane-anesthetized and asleep non-anesthetized neonatal pups, yet the magnitude of activity decreased in the presence of anesthesia (Supplementary Fig. 1). The similarities might be due to the ability of urethane to mimic sleep conditions^28^, the dominant behavioral state of neonatal mice^29^. While dual-hit GE mice have been reported to have profoundly altered network activity and neuronal firing at neonatal age when compared with controls^20^, it is still unclear whether the dysfunction equally affects the local circuitry of all cortical layers. To fill this knowledge gap, we firstly monitored the layer-specific differences between oscillatory patterns of control and dual-hit GE mice. Profound differences in the occurrence, duration and broad band power of oscillatory events were detected when comparing the two groups of mice (Fig. 1, Table 1, Supplementary Fig. 2). However, these detected differences were similar across layers. This might be due on the one hand, to a layer-unspecific overall damping of entrainment in dual-hit GE mice and on the other hand, to non-specific conduction synchrony within a rather small tissue volume (300-400 μm radius). To discriminate between the two sources, in a second step, we investigated the layer-specific firing rate and timing by oscillatory phase, which are not contaminated by non-specific volume conduction^30^. The logarithmic firing rate of neurons in layer II/III of PL in GE mice (0.61 ± 0.04 spikes/s) was significantly (p<0.001) reduced when compared to controls (−2.1 ± 0.1 spikes/s) (Fig. 1e). In contrast, neurons in layer V/VI similarly fired in control (−0.95 ± 0.05 spikes/s) and GE mice (−1.3 ± 0.2 spikes/s). The timing of neuronal firing was also disturbed and lost its precision in layer II/III but not layer V/VI neurons of GE mice when compared to controls (Fig. 1f-h). When averaging over frequency bands the strongest effects (p<0.001) were identified for beta frequency (12-30 Hz) at the level of layer II/III (Fig. 1g), whereas the abnormal spike-LFP synchronization for gamma frequency (30-100 Hz) was less pronounced (p=0.01) (Fig. 1h). In contrast to the weaker timing of firing in layer II/III by beta phase, the firing of layer V/VI neurons in relationship with the beta phase of oscillatory activity was not affected in dual-hit GE mice and the synchrony with the gamma phase was even slightly increased (Fig. 1g,h). On the other hand, the timing of spiking by the theta phase in both layers II/III and V/VI was similar in control and dual-hit GE mice (Supplementary Fig. 2e).

**Fig. 1.**
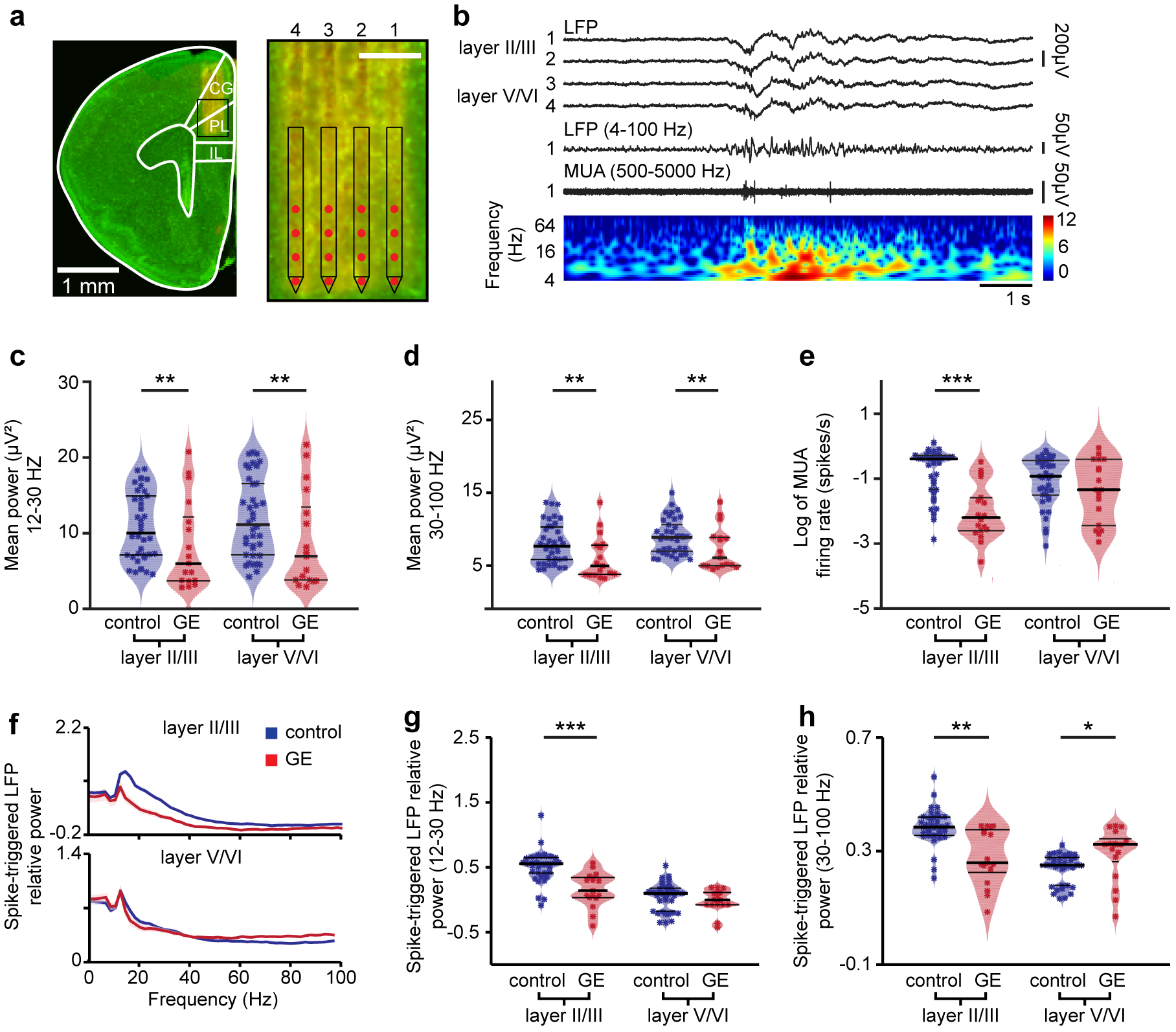
Abnormal patterns of discontinuous oscillatory activity and neuronal firing over the layers of prelimbic cortex of neonatal dual-hit GE mice. (**a**) Digital photomontage reconstructing the position of a 4-shank DiI-labeled recording electrode in the PL of a Nissl-stained 100 μm-thick coronal section (green) from a P9 mouse. Inset, the position of recording sites (red) over the prelimbic layers is displayed at higher magnification. Scale bar, 200 μm. (**b**) Characteristic discontinuous oscillatory activity recorded in layer II/III and V/VI of PL before (top) and after band pass (4-100 Hz) filtering (middle, recording site 1 in layer II/III), and the corresponding MUA after band pass (500-5000 Hz) filtering (bottom, recording site 1 in layer II/III). Color coded frequency plot shows the wavelet of the LFP (recording site 1) at identical time scale. (**c**) Violin plot displaying the power in beta frequency band of oscillations in layer II/III and layer V/VI of the prelimbic cortex of control (blue, n=38) and GE (red, n=18) mice. (**d**) Same as (c) for the power in gamma frequency band. (**e**) Same as (c) for MUA firing rate. (**f**) Plots of frequency-dependent relative power of spike-triggered LFP in layer II/III (top) and V/VI (bottom) of control (blue) and GE (red) mice. (**g**) Violin plot displaying the relative power of spike-triggered LFP in beta band for layer II/III and layer V/VI of control (blue, n=38) and GE (red, n=18) mice. (**h**) Same as (g) for the LFP in gamma band. For (c)-(e), (g), and (h) data is presented as median with 25^th^ and 75^th^ percentile and single data points are shown as asterisks. The shaded area represents the probability distribution of the variable. *P<0.05, **P<0.01 and ***P<0.001, robust ANCOVA with age as covariate (c-e) and yuen’s bootstrap test (g-h) with 20% level of trimming for the mean.

**Table 1.**
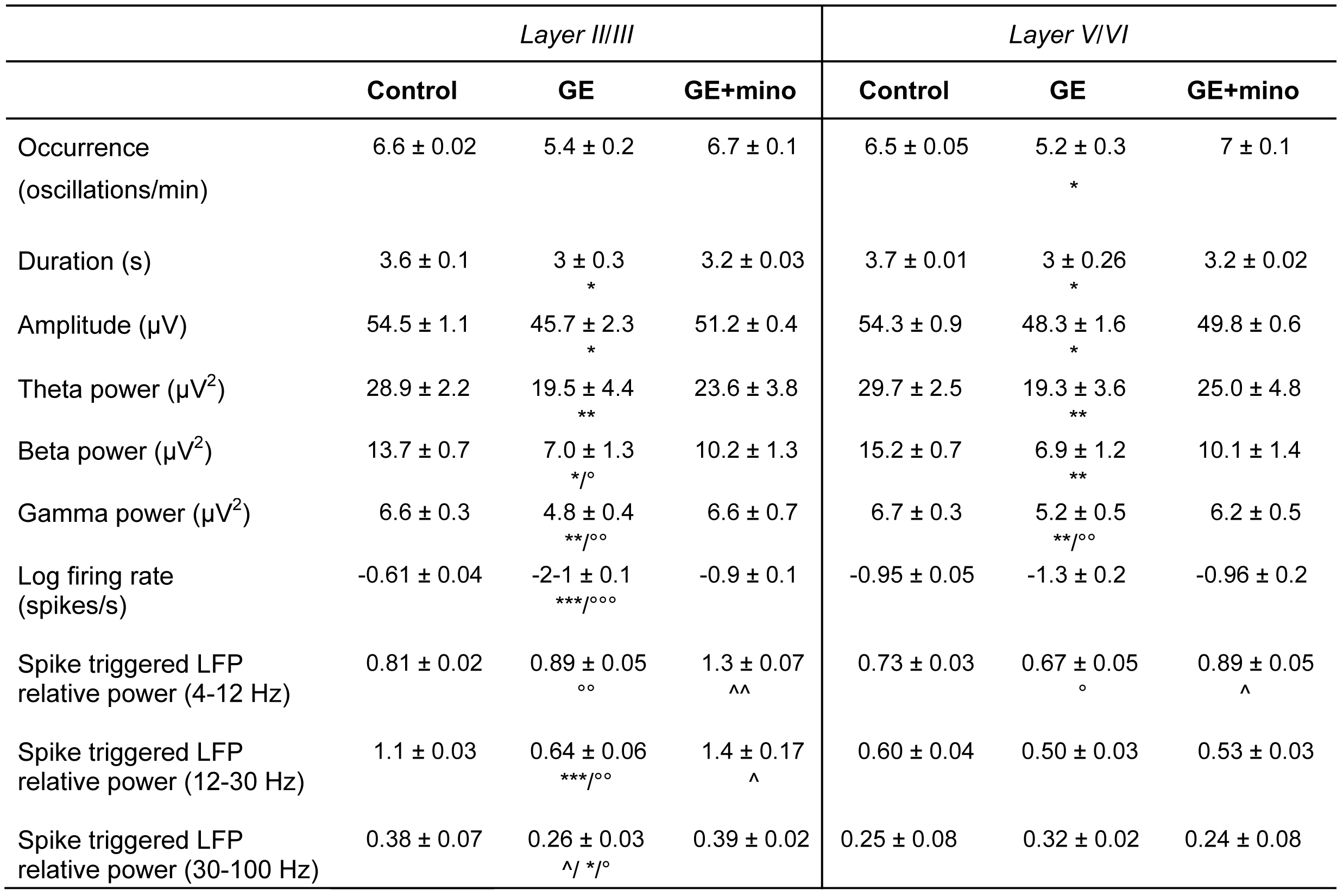
Properties of oscillatory activity and MUA in layer II/III and layer V/VI of PL from control, GE and GE_mino_ mice. Asterisks (*P<0.05, **P<0.01 and ***P<0.001) indicate significance level between control and GE mice. Circles (°P<0.05, °°P<0.01 and °°°P<0.001) indicate significance level between GE and GE_mino_ mice. Carets (ˆP<0.05, ˆˆP<0.01 and ˆˆˆP<0.001) indicate significance level between control and GE_mino_ mice.

These results demonstrate abnormal beta/gamma band oscillations and entrainment of layer II/III of PL in dual-hit GE mice.

### Beta-gamma band dysfunction of prelimbic circuits in dual-hit GE mice results from abnormal activation of layer II/III pyramidal neurons

In developing circuits beta band oscillatory activity has been recently shown to require the activation of pyramidal neurons in layer II/III (PYRs_II/III_) but not V/VI (PYRs_V/VI_) of PL^31^. Therefore, the weaker beta entrainment of prelimbic circuits and coupling of neuronal firing to beta phase identified in GE mice might result from dysfunction of PYRs_II/III_. To test this hypothesis, we monitored the effects of light activation of prelimbic neurons that were transfected with light-sensitive proteins and the red fluorescent protein tDimer2. Using a recently established protocol for optogenetic manipulation of developing circuits^31,32^, we achieved cell-type, layer-, and area-specific transfection of neurons by *in utero* electroporation (IUE) (Supplementary Fig. 3a,b). Constructs coding for the highly efficient fast-kinetics double mutant ChR2 E123 T159 (ET/TC)^33^ were transfected by IUE at embryonic day (E) 15.5 and 12.5 for selective targeting of layer II/III and V/VI, respectively. Staining for NeuN showed that a similar fraction of neurons was transfected in control (34.7 ± 0.8%, n=13 pups) and GE mice (32.0 ± 0.7%, n=8 pups). The pyramidal-like shape and orientation of primary dendrites as well as the expression of CaMKII and the absence of positive staining for GABA confirmed that the expression constructs are exclusively integrated into cell lineages of pyramidal neurons (Supplementary Fig. 3c,d). Omission of ChR2(ET/TC) from the expression construct (i.e. opsin-free) yields similar expression rates and distribution of tDimer2-positive neurons. Moreover, the success rate of transfection by IUE was similar in control and dual-hit GE mice in the presence and absence of opsin (Supplementary Fig. 3e).

The transfection procedure by IUE had no major effects on the overall development of animals (Supplementary Fig. 3f-k). While IUE caused significant reduction of litter size (non-electroporated: 8.3 ± 1.1 pups / litter; IUE: 4.6 ± 1.3 pups / litter; p=0.03), all investigated pups had similar body length, tail length and weight during the early postnatal period. Vibrissa placing, surface righting, and cliff aversion reflexes were also not affected by IUE or transfection of neurons with opsins (Supplementary Fig. 3i-k). These data indicate that the overall somatic development of all investigated pups during embryonic and early postnatal stage is not affected by the transfection procedure.

We firstly assessed the efficiency of light stimulation in inducing action potentials (APs) in prelimbic neurons of control and dual-hit GE mice *in vitro*. For this, whole-cell patch-clamp recordings were performed from tDimer2-positive PYRs_II/III_ (n=42) and PYRs_V/VI_ (n=38) in coronal slices containing the PL from P8-10 mice after IUE at E15.5 and E12.5, respectively. In line with the previously reported “inside-out” pattern of cortical maturation and correspondingly, the more mature profile of neurons in deep vs. superficial layers, PYRs_II/III_ and PYRs_V/VI_ in control mice significantly differed in some of their passive and active membrane properties^31^. However, in dual-hit GE mice the resting membrane potential of PYRs_II/III_ (−53.2 ± 0.37 mV) was more positive when compared with controls (−63.2 ± 0.3mV, p=2*10^−4^) and the halfwidth of action potentials increased (44.8 ± 0.80 mV vs. 29.2 ± 0.36 mV in controls, p=0.018). These alterations of intrinsic neuronal properties point to the immaturity of PYRs_II/III_ in GE Mice. On the contrary, no significant changes in the properties of PYRs_V/VI_ were detected (Supplementary Fig. 4a-d). The passive and active properties of ChR2(ET/TC)-transfected neurons were similar to those previously reported for age-matched mice^31^. Pulsed light stimulation (3 ms, 473 nm, 5.2 mW/mm^2^;) depolarized transfected fluorescently-labeled neurons and led to robust firing in all pups. The probability of triggering APs by pulsed light stimuli decreased with increasing stimulation frequency, yet it differed in its dynamics in control vs. GE mice. Whereas PYRs_II/III_ of control mice were able to reliably follow light stimulations up to 16 Hz, in GE mice they had a significant firing drop already between 8 and 16 Hz (Supplementary Fig. 4e,f). The most prominent differences between control and GE mice were detected for stimulation frequency of 16 Hz. Light stimulation of PYRs_V/VI_ showed a similar decrease of firing probability with augmenting stimulation frequency in control and GE mice.

To elucidate the consequences of abnormal intrinsic firing preference for oscillatory network entrainment, we monitored the effects of light activation of either PYRs_II/III_ or PYRs_V/VI_ *in vivo*. In controls, activation of PYRs_II/III_ selectively drove the neonatal prelimbic networks in beta-gamma frequency range, whereas activation of PYRs_V/VI_ caused non-specific network activation^31^. We reasoned that if PYRs_II/III_ are indeed the cause of the previously demonstrated disruption of beta-gamma activity in the PL of GE mice, then their light stimulation *in vivo* should not be able to specifically induce oscillations in this range. Multi-site recordings of the LFP and MUA were performed from layer II/III of the PL in control and GE mice before, during and after repetitive stimulations with ramp light stimuli. For each pup, the intensity of light stimulation was set to evoke reliable MUA activity and ranged between 20 and 40 mW/mm^2^. The resulting temperature increase (>0.2 °C) and tissue heating during illumination estimated according to a previously developed model^34^ are below those that have been reported to augment the neuronal firing^35^.

Light stimulation increased the neuronal firing of ChR2(ET/TC)-transfected PYRs_II/III_ and PYRs_V/VI_ in control and GE mice but not of neurons transfected with opsin-free constructs (Fig. 2a-d; 3a-d; Supplementary Fig. 5). The spike discharge was initiated once the power exceeded a certain threshold. For some neurons, the firing decreased towards the end of the ramp stimulations, indicating that, similar to the *in vitro* conditions, their membrane potential reached a depolarizing plateau preventing further spiking. However, for the majority of neurons, the firing rate after stimulus remained higher than before the stimulus (Fig. 2a,b; Fig. 3a,b), suggesting that short-term plasticity or global network activation have been induced by light stimulation in developing circuits. Ramp stimulation revealed major differences in the firing of prelimbic neurons from control and GE mice. While PYRs_II/III_ in controls did not fire randomly during stimulus but, as previously demonstrated, had a preferred inter-spike interval of ∼60 ms, equivalent to a population firing at 16.7 Hz (Fig. 2a,c), a coordinated discharge pattern was absent in all investigated GE mice upon ramp stimulation of PYRs_II/III_ (Fig. 2b,d). In contrast, the firing dynamics of PYRs_V/VI_ was similar in control and GE mice and showed no frequency-specific concentration of firing during ramp stimulation (Fig. 3a,b).

**Fig. 2.**
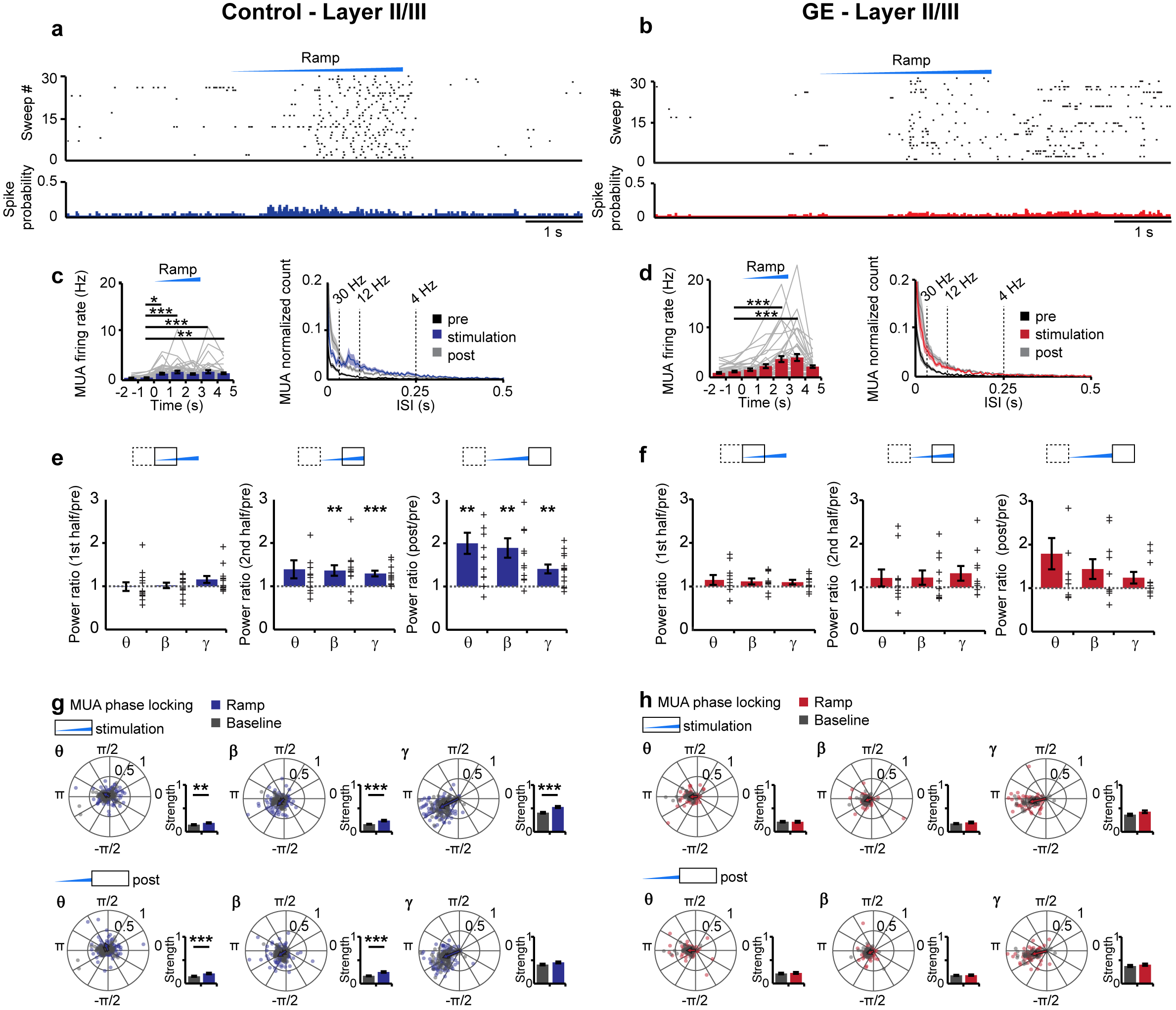
Optogenetic activation of PYRs_II/III_ in control and dual-hit GE mice *in vivo*. (**a**) Representative raster plot and corresponding spike probability histogram displaying the firing of a PYR_II/III_ from a control mouse in response to 30 sweeps of ramp stimulation (473 nm, 3s). (**b**) Same as (a) for transfected PYRs_II/III_ from GE mice. (**c**) Left, bar diagram displaying the mean MUA firing rate in transfected PYRs_II/III_ of control mice in response to ramp illumination. Right, occurrence rate of interspike intervals averaged for 3 s before light stimulation (pre, black), 3 s during ramp stimulation (stimulation, blue) and 3 s after light stimulation (post, gray, n=43 recording sites from 13 pups). (**d**) Same as (c) for transfected PYRs_II_/_III_ from GE mice (red, n=10 pups; MUA of n=40 recording sites). (**e**) Bar diagrams displaying the LFP power for control mice during the first half (1.5 s), the second half (1.5 s) and after (post, 1.5 s) ramp stimulation, normalized to the power before stimulation (pre, 1.5 s). Network activity in theta (9, 4-12 Hz), beta (**(3**, 12-30 Hz) and gamma (y, 30-100 Hz) frequency bands (blue, n=13 pups) is considered. (**f**) Same as (e) for GE mice (red, n=10 pups). (**g**) Polar plots displaying the phase-locking of light-triggered (blue, stimulation 3 s, post 3 s) and spontaneous (gray) MUA to oscillatory activity in PYRs_II_/_III_ of control mice (n=43 recording sites from 13 pups). Bar diagrams display the locking strength. (**h**) Same as (g) for GE mice (red, n=10 pups; MUA of n=40 recording sites). In (c) and (d) gray lines correspond to firing of individual neurons. In (e) and (f) individual values correspond to pups and are displayed as gray crosses. In (g) and (h) individual values from individual units are shown as dots and gray crosses, whereas the arrows correspond to the mean resulting group vectors. Data are presented as mean ± s.e.m. *P<0.05, **P<0.01 and ***P<0.001), one-way repeated-measures analysis of variance (ANOVA) with Bonferroni-corrected *post hoc* analysis (c,d), two-sided t-tests and circular statistic toolbox (e-h).

**Fig. 3.**
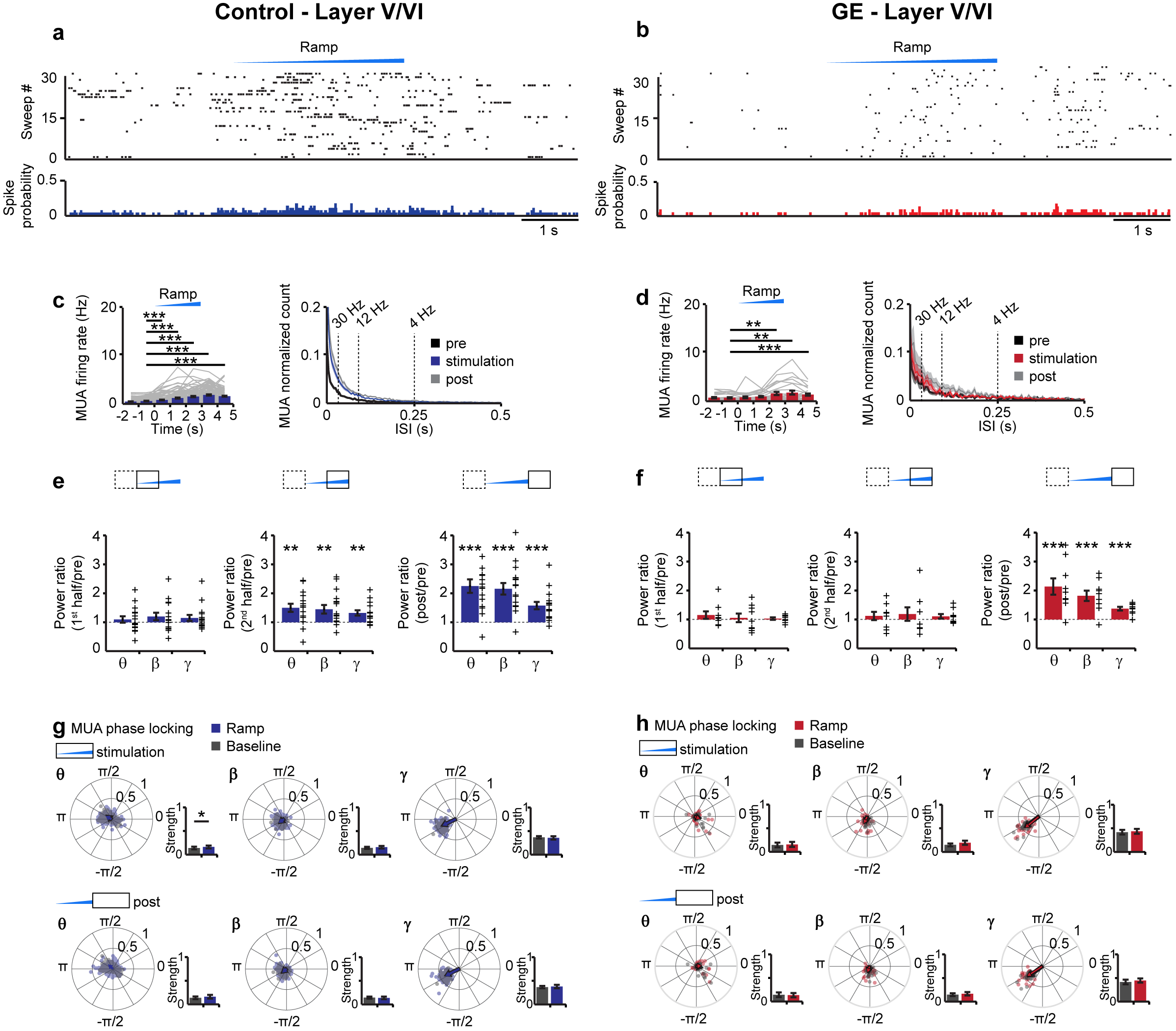
Optogenetic activation of PYRs_V/V_I in control and dual-hit GE mice *in vivo*. (**a**) Representative raster plot and corresponding spike probability histogram displaying the firing of a PYR_V/VI_ from a control mouse in response to 30 sweeps of ramp stimulation (473 nm, 3s). (**b**) Same as (a) for transfected PYRs_V/V_I from GE mice. (**c**) Left, bar diagram displaying the mean MUA firing rate in transfected PYRs_V/V_I of control mice in response to ramp illumination. Right, occurrence rate of interspike intervals averaged for 3 s before light stimulation (pre, black), 3 s during ramp stimulation (stimulation, blue) and 3 s after light stimulation (post, gray, n=117 recording sites from 16 pups). (**d**) Same as (c) for transfected PYRs_V/VI_ from GE mice (red, n=6 pups; MUA of n=27 recording sites). (**e**) Bar diagrams displaying the LFP power for control mice during the first half (1.5 s), the second half (1.5 s) and after (post, 1.5 s) ramp stimulation, normalized to the power before stimulation (pre, 1.5 s). Network activity in theta (9, 4-12 Hz), beta (**(3**, 12-30 Hz) and gamma (y, 30-100 Hz) frequency bands (blue, n=16 pups) is considered. (**f**) Same as (e) for GE mice (red, n=9 pups). (**g**) Polar plots displaying the phase-locking of light-triggered (blue, stimulation 3 s, post 3 s) and spontaneous (grey) MUA to oscillatory activity in PYRs_V/VI_ of control mice (n=117 recording sites from 16 pups). Bar diagrams display the locking strength. (**h**) Same as (g) for GE mice (red, n=6 pups; MUA of n=27 recording sites). In (c,d) gray lines correspond to firing of individual neurons. In (e and f) individual values correspond to pups and are displayed as gray crosses. In (g) and (h) individual values from individual units are shown as dots and gray crosses, whereas the arrows correspond to the mean resulting group vectors. Data are presented as mean ± s.e.m. *P<0.05, **P<0.01 and ***P<0.001), one-way repeated-measures analysis of variance (ANOVA) with Bonferroni-corrected *post hoc* analysis (c,d), two-sided t-tests and circular statistic toolbox (e-h).

To causally prove the contribution of abnormal firing of PYRs_II/III_ to the weaker beta-gamma band entrainment previously identified in the PL of dual-hit GE mice, we tested the effects of ramp stimulations on the discontinuous patterns of oscillatory activity. When compared with pulsed stimulations, ramp stimulations have the advantage of not inducing power contamination by repetitive and fast large-amplitude voltage deflections resulting from simultaneous opening of light-activated channels^31,36^ and to trigger more physiological and not artificially synchronous firing patterns. In control mice, the LFP power in beta and gamma frequency range significantly increased during ramp stimulation of PYRs_II/III_, whereas the theta band activity remained unaffected. Even after the stimulus, the augmented network power persisted, yet lacked frequency specificity (Fig. 2e). In contrast, PYRs_II/III_ in GE mice lost their ability to boost neonatal prelimbic oscillations in frequency-specific manner, since ramp stimulations did not affect LFP power (Fig. 2f). Not only the light-induced inter-spike interval and power of network oscillations were disrupted in GE mice, but also the timing of firing by the oscillatory phase was impaired. We calculated the phase locking of MUA and clustered single-unit activity (SUA) to oscillations for layer II/III in the PL of control and GE mice. In control mice, the neuronal firing was more strongly phase-locked to theta (baseline: 0.15 ± 0.01; ramp: 0.19 ± 0.01; p=0.016), beta (baseline: 0.16 ± 0.01; ramp: 0.24 ± 0.01; p<0.001), and gamma oscillations (baseline: 0.40 ± 0.01; ramp: 0.53 ± 0.01; p<0.001) during ramp stimulation when compared to spontaneous activity (Fig. 2g). In contrast, the phase locking to theta (baseline: 0.20 ± 0.01; ramp: 0.21 ± 0.01; P>0.05), beta (baseline: 0.17 ± 0.01; ramp: 0.19 ± 0.01; P>0.05), and gamma oscillations (baseline: 0.38 ± 0.02; ramp: 0.43 ± 0.03; P>0.05) during stimulation of PYRs_II/III_ in GE mice was similar to the coupling during spontaneous activity (Fig. 2h).

In line with the frequency-unspecific augmentation of firing rate after light activation of PYRs_V/VI_ in control mice, the LFP power in all frequency bands increased during stimulation and remained at a high level even after it (Fig. 3e). In GE mice the power augmentation started only after the stimulus (Fig. 3f). Neither in control nor in GE mice, light activation of PYRs_V/VI_ strengthened the timing of neuronal firing by the phase of oscillatory activity (Fig. 3g,h).

Thus, the reduced beta-gamma oscillatory activity in the PL of neonatal dual-hit GE mice results from dysfunction of firing dynamics of PYRs_II/III_.

### Layer II/III pyramidal neurons of neonatal dual-hit GE mice show major morphological and synaptic deficits

The selective dysfunction of PYRs_II/III_ and the related abnormal network activity in GE mice might be related to abnormal morphology and connectivity of these neurons at neonatal age. To test this hypothesis, we undertook a detailed histological examination of the cytoarchitecture of tDimer-labeled pyramidal neurons in layers II/III and V/VI of P10 control and GE mice. PYRs_II/III_ of GE mice but not PYRs_V/VI_, showed a significant reduction in the soma size when compared to neurons of controls (n=21 neurons for every condition; condition effect, p=0.04 and p>0.05, respectively). The complexity of dendritic branching was assessed by Sholl analysis of three-dimensionally reconstructed PYRs_II/III_ and PYRs_V/VI_. When compared to controls, PYRs_II/III_ of GE mice had major reduction in dendritic branching (condition effect, p=2*10^−16^) (Fig. 4a-c). These deficits were particularly prominent within a radius of 25-115 μm from the cell soma center (p<0.05 for all pairwise comparisons). In accordance with our electrophysiological results, we found no significant differences in the complexity of dendritic arborization for PYRs_V/VI_ of GE and control mice (Fig. 4d-f).

**Fig. 4.**
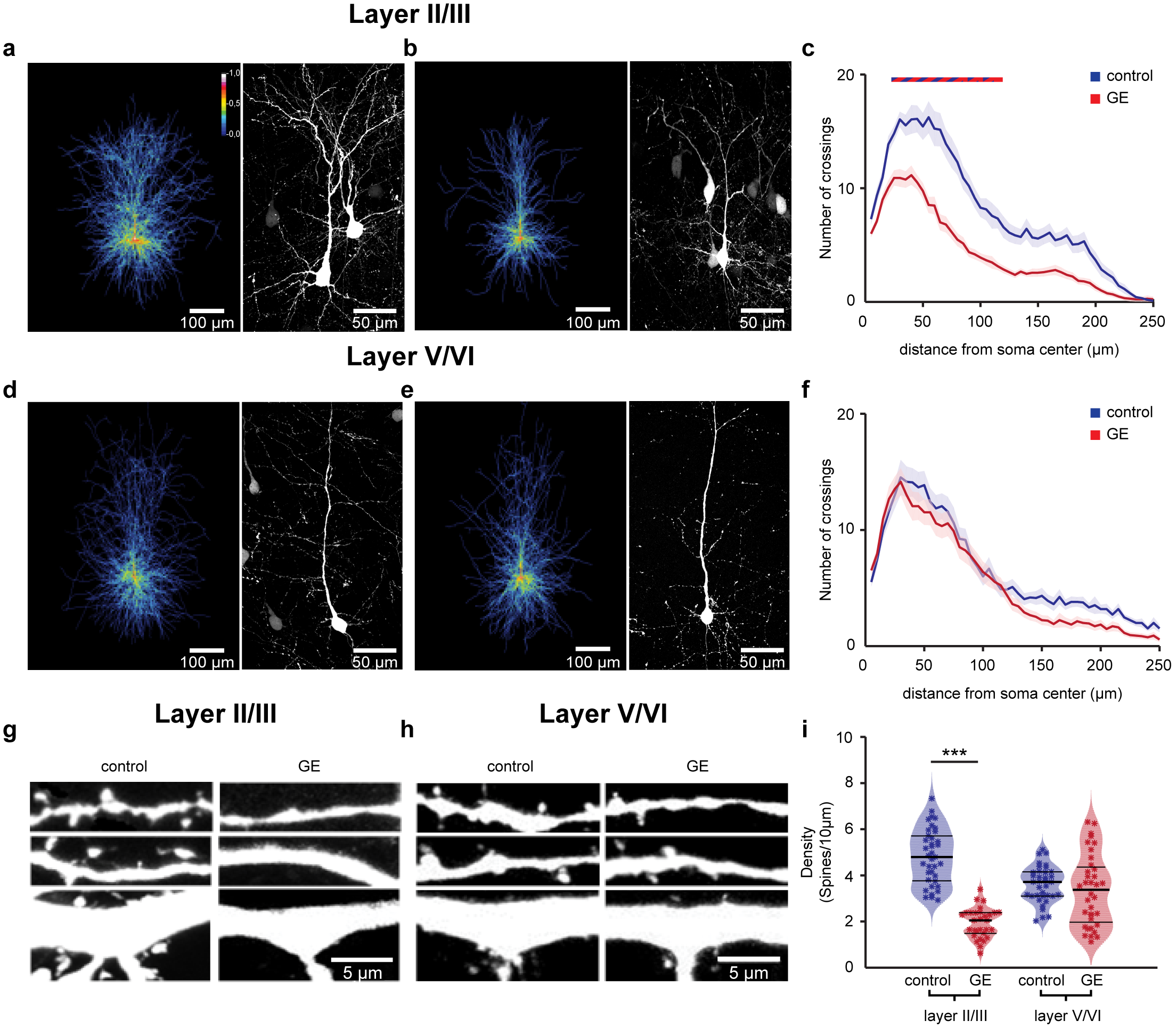
Simplified dendritic arborization and reduced spine density in PYRs_II/III_ of dual-hit GE mice. (**a**) Left, heatmap displaying an overlay of all traced dendrites of transfected PYRs_II/III_ in control mice. Right, photograph of a representative PYRs_II/III_ in a P10 mouse. (**b**) Same as (a) for a P10 duaö-hit GE mouse. (**c**) Graph displaying the average number of dendritic intersections within a 250 μm radius from the soma center of PYRs_II/III_ in control (blue, n=21 neurons from 3 pups) and GE (red, n=21 neurons from 3 pups) mice. Blue/red bar indicates significant difference between control and GE mice. (**d-f**) Same as (a-c) for PYRs_V/VI_ from control (blue, n=21 neurons from 3 pups) and GE (red, n=21 neurons from 3 pups) mice. (**g**) Photograph displays representative basal (top), secondary apical (middle), proximal oblique and apical (bottom) dendrites of a PYR_II/III_ from a P10 control mouse (left) and a P10 GE mouse (right). (**h**) Same as (g) for PYRs_V/VI_. (**i**) Violin plot displaying the average spine concentration on dendrites from PYRs_II/III_ of control (blue, n=39 dendrites from 13 neurons) and GE (red, n=30 dendrites from 10 neurons) mice. In (c) and (f) data is presented as mean ± s.e.m. In (i) data is presented as median with 25^th^ and 75^th^ percentile, and single data points are displayed as asterisks. The shaded area represents the probability distribution of the variable. *P<0.05, **P<0.01 and ***P<0.001, linear mixed-effect model.

Next, we examined the spine density along the dendrites of PYRs_II/III_ and PYRs_V/VI_ whose dendritic morphology we had previously analyzed. PYRs_II/III_ of GE mice (n=10 neurons) has significantly lower density when compared to controls (n=13 neurons, condition effect, p=7*10^−4^), whereas the values were comparable for PYRs_V/VI_ of control (n=9 neurons) and GE mice (n=9 neurons, condition effect, p>0.05) (Fig. 4g-i). The magnitude of density reduction was similar for different types of dendrites (apical and proximal oblique dendrites, secondary apical dendrites, and basal dendrites; condition effect p<0.001) (Supplementary Fig. 6).

The simplified dendritic arborization and the decreased spine density of PYRs_II/III_, but not PYRs_V/VI_ further confirm the layer-specific dysfunction in dual-hit GE mice.

### Blocking the microglia inflammatory response shortly after birth rescues the prelimbic pyramidal firing and oscillatory entrainment in dual-hit GE mice

We next set out to determine whether the morphological and functional deficits of PYRs_II/III_ in the PL of GE mice could be rescued during early development. Microglia have been reported to sculpt the developing circuits by engulfing and remodeling synapses in an activity-dependent manner^37,38^. Transient perturbations in the development of microglia, such as those induced by the used environmental stressor, maternal immune activation (MIA), have far reaching effects on adult neuronal function and behavior^39,40^ that have been linked to mental illness ^41^. Moreover, mutations in the C4 gene, a complement component that is required for synapse elimination by microglia^38^, has been linked to schizophrenia^41^.

In accordance with this stream of evidence, investigation of microglia in the PL of dual-hit GE mice revealed major alterations in this population of cells in GE mice. When compared with controls, not only was microglia number significantly augmented (+47%, p=3*10^−6^), but also morphological features such as area and cell-spread were likewise significantly increased by 29% (p=4*10^−7^) and 25% (p=8*10^−14^), respectively (Fig. 5). Moreover, microglia cell perimeter and roundness, but not eccentricity, were also substantially changed in dual-hit GE mice (Supplementary Fig. 7). Such morphological abnormalities are suggestive of a hyper-mature microglial population, as cell size and arborization increase through development^42^. Our results are in line with a recent report indicating that, at a similar time point to the one that we investigated, MIA induces a transcriptomic profile reminiscent of adult microglial cells^39^.

**Fig. 5.**
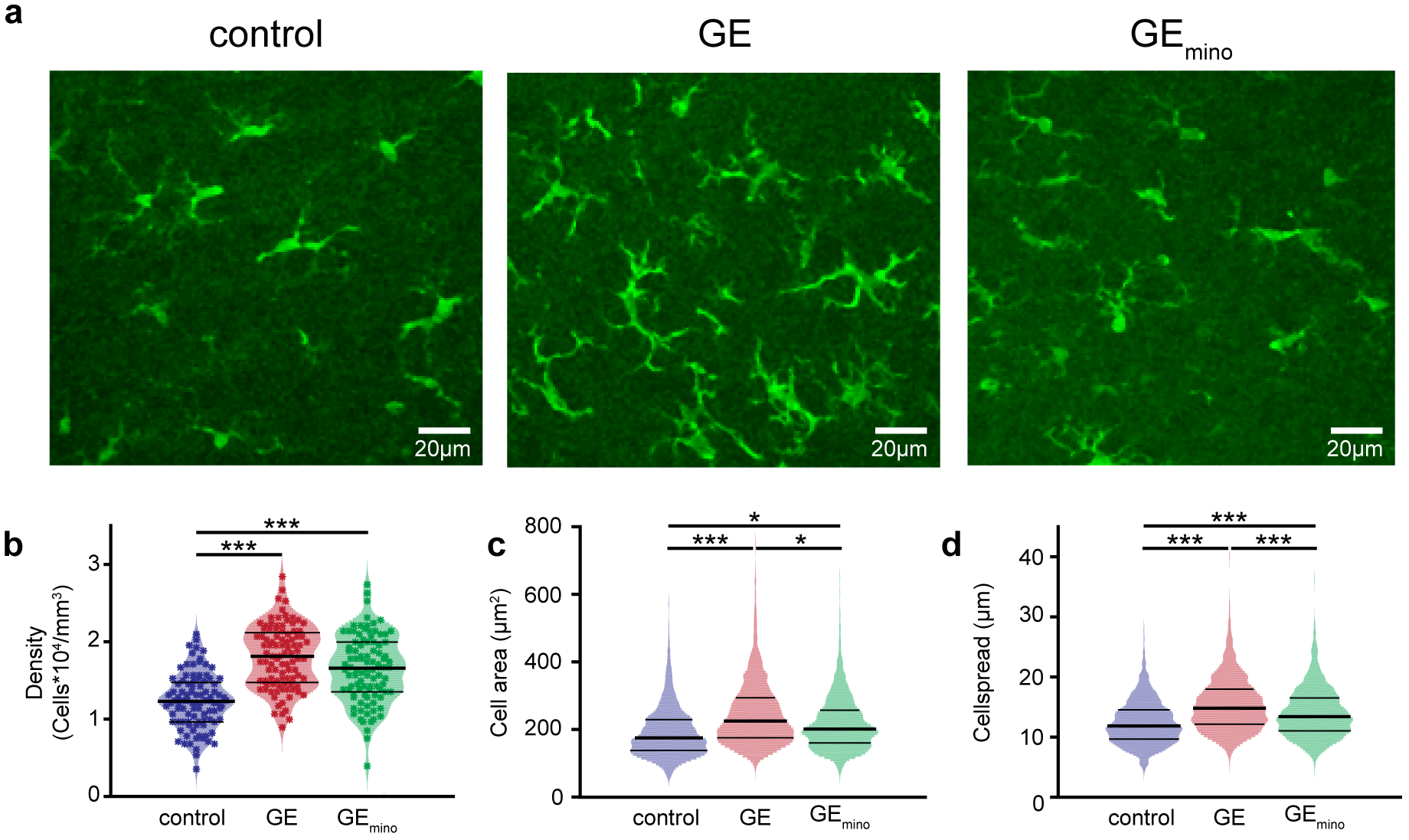
Altered microglial cell population in dual-hit GE mice is partially restored by minocycline treatment. (**a**) Photographs of Iba−1-stained microglial cells in the PL of a P10 control mouse (left), of a P10 GE mouse (center) and of a P10 GE_mino_ mouse (right). (**b**) Violin plot displaying the average density of Iba-1-stained cells in the PL of control (blue, n=64 images from 4 pups), GE (red, n=64 images from 4 pups) and GE_mino_ mice (green, n=64 images from 4 pups). (**c**) Same as (b) for cell area. (**d**) Same as (b) for cellspread. For (c) and (d), n=1250, 1738 and 1614 cells, respectively, from 12 sections of 4 pups, for all three conditions. Data is presented as median with 25^th^ and 75^th^ percentile. In (b) single data points are displayed as asterisks, whereas in (c) and (d) single data points are omitted due to their high number. The shaded area represents the probability distribution of the variable. *P<0.05, **P<0.01 and ***P<0.001, linear mixed-effect model.

To rescue the hyper-maturation of microglia, we exposed GE mice during neonatal development (from P1 to P8) to minocycline, a tetracycline antibiotic that blocks the stress-induced inflammatory responses of these cells^43^. While minocycline-treated GE (GE_mino_) mice had only a weak and non-significant reduction in the number of microglial cells (−13%, p=0.17), microglia showed a reduced area (−35%, p=0.015) and cellspread (−11%, p=8*10^−4^). These data indicate that minocycline treatment partially restores the microglia alterations in GE mice.

To prove the contribution of microglial deficits to the dysfunction of PYRs_II/III_, we assessed the morphology and function of these neurons after minocycline treatment. First, Sholl analysis of three-dimensionally reconstructed tDimer-positive PYRs_II/III_ (n=21) from GE_mino_ mice showed that the magnitude of dendritic branching was fully restored after treatment, being similar to that of controls (condition effect, p>0.05) (Fig. 6a,b). Minocycline treatment restored the synaptic deficits too. PYRs_II/III_ from GE_mino_ mice (n=12 neurons) had a similar spine density as those from control mice (condition effect, p>0.05) and significantly increased when compared to GE mice (condition effect, p=5*10^−5^) (Fig. 6c,d). The effect was similar across the different types of dendrites that were analyzed (apical and proximal oblique dendrites, secondary apical dendrites, and basal dendrites; condition effect >0.05 and <0.001, when compared to neurons from control and GE mice, respectively).

**Fig. 6.**
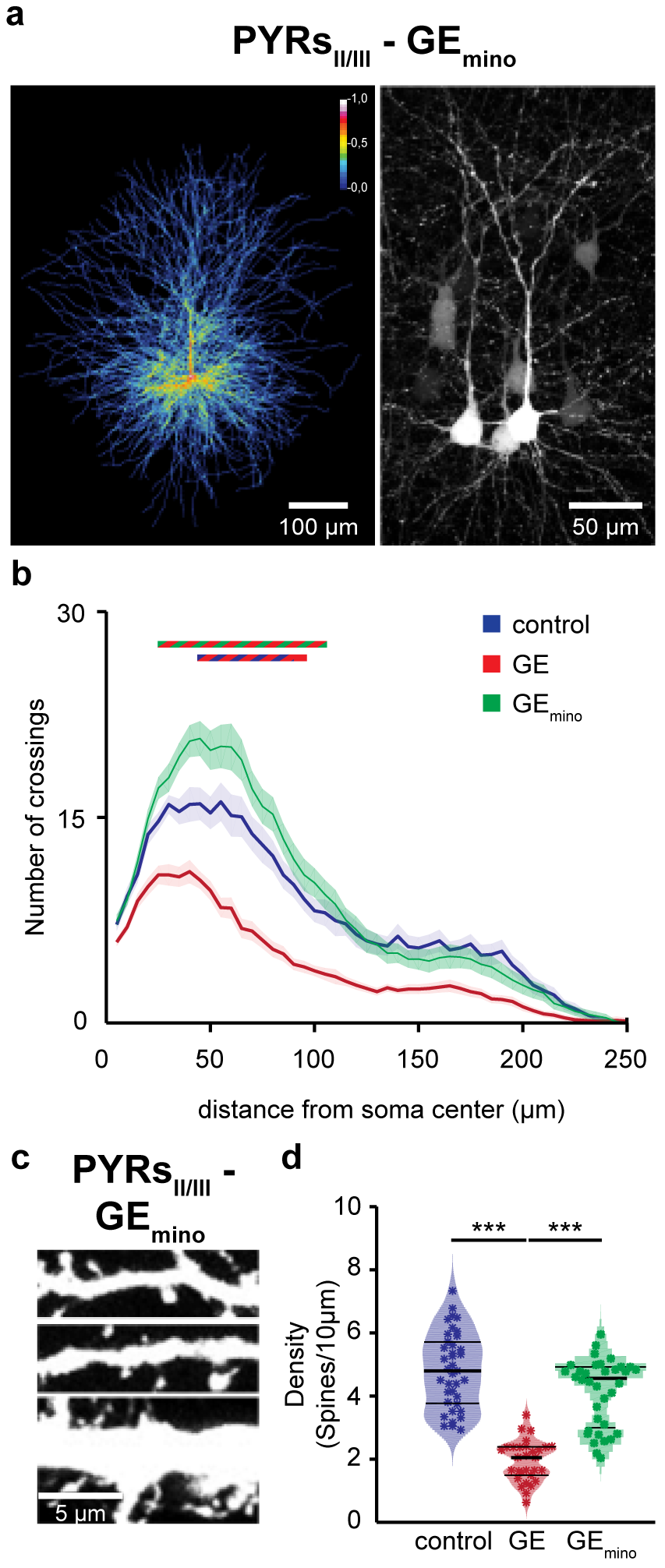
Minocycline treatment rescues the abnormal structure of PYRs_II/III_ in GE mice. (**a**) Left, heatmap displaying an overlay of all traced dendrites of transfected PYRs_II/III_ in GE_mino_ mice. Right, photograph of a representative PYR_II/III_ in a P10 GE_mino_ mouse. (**b**) Graph displaying the average number of dendritic intersections within a 250 μm radius from the soma center of PYRs_II/III_ in control (blue, n=21 neurons from 3 pups), GE (red, n=21 neurons from 3 pups), and GE_mino_ (green, n=21 neurons from 3 pups) mice. Blue/red and green/red bars indicate significant difference between control and GE mice and GE and GE_mino_ mice, respectively. (**c**) Photograph displays representative basal (top), secondary apical (middle), proximal oblique and apical (bottom) dendrites of a PYR_II/III_ from a P10 GE_mino_ mouse. (**d**) Violin plot displaying the average spine concentration on dendrites from PYRs_II/III_ of control (blue, n=39 dendrites from 13 neurons), GE (red, n=30 dendrites from 10 neurons) and GE_mino_ (green, n=36 dendrites from 12 neurons) mice. In (b) data is presented as mean ± s.e.m. In (d) data is presented as median with 25^th^ and 75^th^ percentile and single data points are displayed as asterisks. The shaded area represents the probability distribution of the variable. ***P<0.001, linear mixed-effect model.

Second, we assessed the properties of prelimbic network oscillations and neuronal firing in GE_mino_ mice, and compared them with those from control and GE mice. The power in beta and gamma band of prelimbic oscillations recorded in layer II/III were similar in control and minocycline-treated GE mice (Fig. 7a,b, Table 1). Similarly, the prelimbic firing rate and timing by oscillatory phase were rescued by restoring the microglial network (Fig. 7c-f). The firing rate of neurons in layer II/III was similar for controls (log values −0.61 ± 0.04) and GE_mino_ (log values −0.9 ± 0.1, p>0.05). The timing of prelimbic firing in layers II/III of GE_mino_ mice, as measured by spike-triggered LFP power, was not only rescued, but even slightly increased when compared to controls for both beta (p=0.049) and gamma (p=0.049) band (Fig. 7d-f). In contrast to the profound changes observed in layer II/III after minocycline treatment, the network activity and neuronal firing in layer V/VI of PL from GE_mino_ mice remained largely unaffected (Supplementary Fig. 8).

**Fig. 7.**
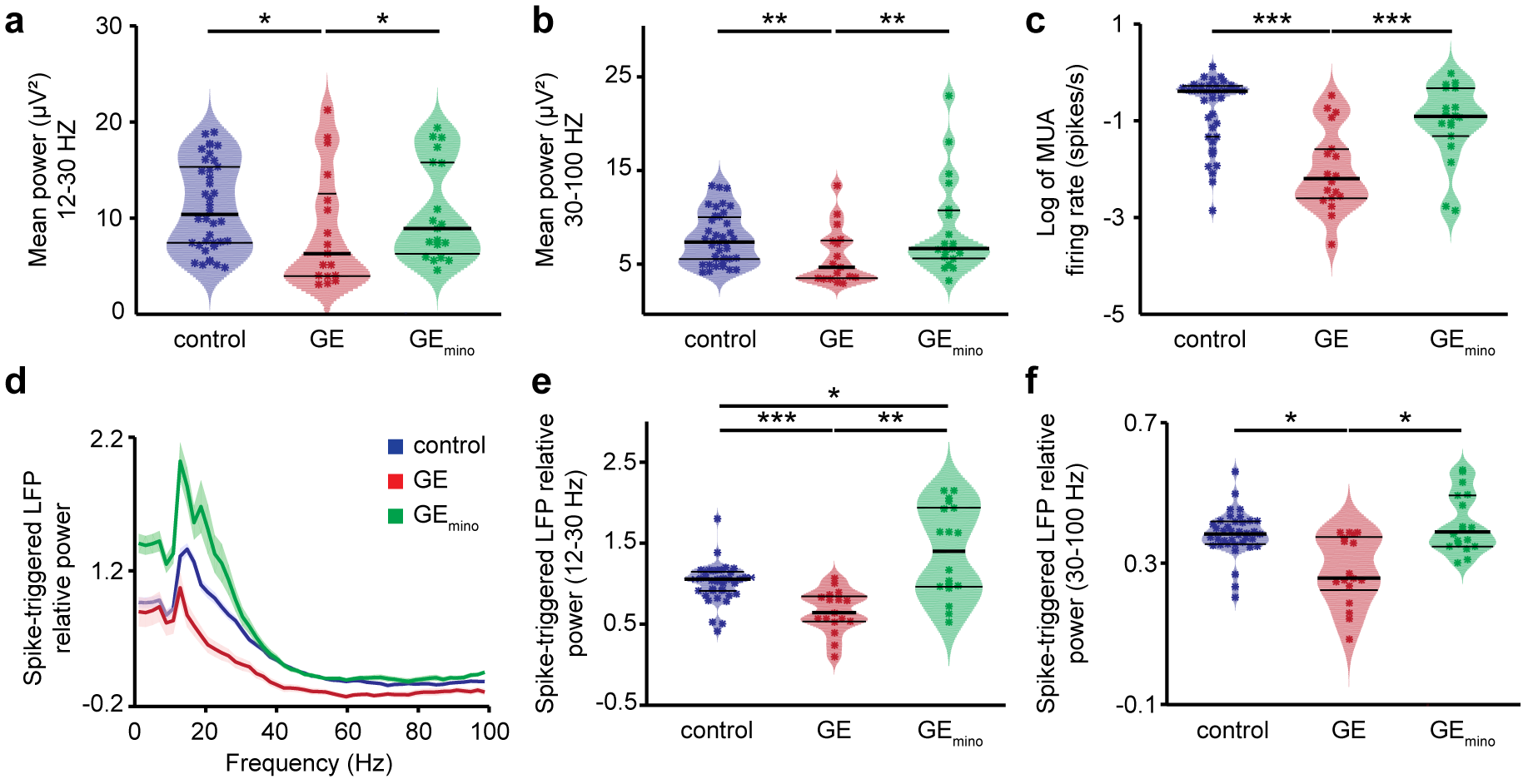
Minocycline treatment rescues core dysfunctions in prelimbic layer II/III of neonatal dual-hit GE mice. (**a**) Violin plot displaying the beta band power of oscillations in layer II/III of the PL of control (blue, n=38), GE (red, n=18), and GE_mino_ (green, n=18) mice. (**b**) Same as (a) for the gamma power. (**c**). Same as (A) for MUA firing rate. (**d**) Plots of frequency-dependent relative power of spike-triggered LFP in layer II/III (top) of control (blue), GE (red), and GE_mino_ (green) mice. (**e**) Violin plot displaying the relative power of spike-triggered LFP in beta band for layer II/III of control (blue, n=38), GE (red, n=18), and GE_mino_ (green, n=18) mice. (**f**) Same as (e) for the LFP in gamma band. In (d) data is presented as mean ± sem. In (a)-(c), (e), and (f) data is presented as median with 25^th^ and 75^th^ percentile and single data points are shown as asterisks. The shaded area represents the probability distribution of the variable. *P<0.05, **P<0.01 and ***P<0.001, robust ANCOVA with age as covariate and 20% level of trimming for the mean (a)-(c) and robust ANOVA with 20% level of trimming for the mean (e), (f), with Bonferroni-Holm *post-hoc* analysis.

These data indicate that the abnormal firing and network coupling patterns in the PL of dual-hit GE mice can be rescued by inhibition of inflammatory response.

## DISCUSSION

While neurodevelopmental miswiring has been postulated to result in major functional and behavioral deficits at adulthood, the mechanisms of early impairment are still largely unresolved. A major consequence of this knowledge gap is the poor understanding of disease pathophysiology that hampers the development of tailored therapies for mental illness. Towards the aim of elucidating the substrate of developmental dysfunction, the present study uncovers layer- and cell type-specific deficits in the PL of neonatal mice reproducing the gene-environment interactions involved in the pathogenesis of psychiatric disorders. Combining *in vivo* electrophysiology and optogenetics with in-depth morphological analysis and pharmacology we showed that in dual-hit GE mice (i) the lower entrainment of neonatal prefrontal circuits in beta-gamma oscillations results from structural and functional deficits of layer II/III pyramidal neurons; (ii) pharmacological intervention on hyper-mature microglia cells restores the morphology, synaptic function, firing and oscillatory patterns in the local prefrontal circuits. These findings highlight the major contribution of glutamatergic dysfunction to the abnormal refinement of circuits during development and support the hypothesis that the excitation/inhibition imbalance in the context of mental diseases emerges already at neonatal age.

### Wiring of prefrontal circuits at neonatal age: check point of cognitive maturation

Anatomical investigations revealed that, while the PFC develops according to a similar time schedule as do other neocortical areas, such as primary cortices, some maturation events (e.g. volumetric decline, growth and pruning of afferents and efferents) are protracted^44^. Correspondingly, the prefrontal patterns of coordinated activity share the general spatial and temporal organization of early neocortical oscillations^45^, yet they emerge later and have a frequency-specific structure. In rodents, the discontinuous oscillatory activity of PFC appears 1-2 days later than in the hippocampus and 2-3 days later than in primary visual and somatosensory cortices^5,46,47^. The neonatal oscillations are detectable and have similar organization in both urethane-anesthetized and non-anesthetized rats and mice, yet their magnitude decreased under anesthesia, as shown by the present and previous studies^27^.

In line with the strong long-range connections identified in adults^48,49^, the rodent PFC receives a strong glutamatergic drive from the hippocampus that mainly targets deep layers towards the end of the 1^st^ postnatal week^5,20,50^. At functional level, this connectivity enables that the hippocampal bursts in theta frequency band entrain prefrontal circuits that generate fast oscillations superimposed on the theta ground rhythm relayed from the hippocampus. These nested-gamma spindle bursts seem to be unique to the developing PFC and reflect the wiring between prefrontal layers. Layer V/VI neurons are tightly interconnected with pyramidal neurons in layer II/III. In line with our present and previous data pyramidal neurons in layer II/III are the generators of beta-low gamma oscillations, whereas pyramidal neurons in layer V/VI contribute to the overall activation of PFC, gatekeeping long-range inputs, such as from hippocampus^27^.

These findings show that frequency-specific communication within local prefrontal circuits and the directed coupling within prefrontal-hippocampal networks emerge very early. Such precise interactions might represent the pre-requisite for functional entrainment of adult networks and cognitive performance. Abundant literature highlighted the link between theta band hippocampal activation and fast oscillatory entrainment of prefrontal circuits during various cognitive tasks at adulthood^51–55^. Currently, only few attempts have been made to directly prove the role of timed interactions during development for the later emergence of network function and adult behavior. Recently, the key role of vasoactive intestinal peptide (VIP) and cholecystokinine (CCK)-positive interneurons for cortical circuit development has been demonstrated^56,57^. In the same line, our findings identify pyramidal neurons in layer II/III as generators of early activity, facilitating the coupling within local neocortical circuits and glutamatergic communication between upper and deeper layers^58^. Identification of key cellular elements controlling circuit development opens new perspectives for the interrogation of long-term network effects.

### Mechanisms of abnormal wiring in prefrontal circuits of neonatal dual-hit GE mice

The pathogenesis of cognitive dysfunction in major psychiatric disorders has been reported to involve interactions between a large number of susceptibility genes and environmental factors that might act at diverse stages of development^59,60^. In the present study, we used a dual-hit mouse model that mimics the abnormally translocated DISC1 gene^61–63^, on the one hand, and the *in utero* viral infection causing maternal immune activation, on the other hand^64–66^. We chose this combination because previous studies documented their clear link to human pathology^67^. While each of the factors (i.e. either genetic or environmental) leads to structural, functional and cognitive deficits of weak to moderate magnitude^21^, it is only their combination that has been found to produce a neurobehavioral phenotype at adulthood that resembles aspects of mental illness. The used dual-hit GE mice recapitulate this feature, as our previous investigation revealed that the prefrontal-hippocampal networks accounting for mnemonic and executive abilities at adult age abnormally mature only when both hits co-occur, whereas single-hit models show a largely normal network development^20^. The patterns of coordinated activity in PFC and hippocampus appeared disorganized and the long-range coupling between them was weaker at neonatal age. In contrast, at pre-juvenile age, when the first memory abilities arise, an exaggerated activation and coupling has been detected within prefrontal-hippocampal networks.

While these data experimentally demonstrate the developmental origin of dysfunction in dual-hit GE mice, they do not mechanistically explain the network and behavioral deficits. They might result from either abnormal maturation of local prefrontal networks, a weaker theta activity in hippocampus or sparser connectivity between the two areas. The present results fill the knowledge gap and identify the pyramidal neurons of layer II/III as key players of developmental miswiring, whereas pyramidal neurons in deeper cortical layers are indistinguishable in their structure and function in controls and dual-hit GE mice. The disorganized patterns of oscillatory activity in the PL result from layer II/III neurons that lost their timed firing and cannot generate the entrainment of local circuits in beta-low gamma frequencies. On its turn, the neuronal spiking is controlled by the inputs that these neurons receive. Taking into account the oversimplified dendritic arborization and reduced number of spines it is likely that pyramidal neurons in layer II/III of PL from dual-hit GE mice receive less inputs, which are randomly timed. The cell type and layer-dependent structural abnormalities in PFC (e.g. decreased dendritic spine density) have also been detected in schizophrenia patients and related to abnormalities in the excitatory transmission^68,69^. Despite the integrity of pyramidal neurons in layers V/VI both at morphological and functional level, the local prefrontal circuitry relying on dense vertical and horizontal interactions between upper and deeper layers^70^ is compromised in dual-hit GE mice. Therefore, the theta hippocampal drive targeting layer V/VI pyramidal neurons cannot optimally entrain the PFC.

The selective dysfunction of pyramidal neurons in prefrontal layer II/III was rescued by inhibiting the inflammatory response and restoring the function of microglia in dual-hit GE mice. Abundant literature documented that microglial processes dynamically and preferentially interact with the dendritic spines of pyramidal neurons^37,71,72^. Altered number of activated microglia has been found in the brain of MIA offspring^73,74^. Resulting from maternal infection, the stimulation of cytokine pathways^65^ and the elevated glucocorticoid concentrations ^75^ have been proposed to interfere with developmental processes, such as neuronal proliferation, differentiation, and synaptogenesis^76^. Moreover, overactivation of microglia during development seems to be sufficient to induce persistent changes in glutamatergic neurotransmission^77^, yet the concept still needs solid experimental evidence. During the first postnatal week, these MIA-induced deficits alone seem to have no functional readout, since neonatal one-hit environmental mice (i.e MIA offspring) have largely normal firing and network activity patterns^20^. Solely the combination with genetic risk factors, such as mutant DISC1, causes microglia-dependent early circuit miswiring, as reported in the present study. DISC1 has a key role in neuronal proliferation and migration as well as in development and maintenance of synapses. However, the phenotypes in the mouse models of mutated DISC1 are rather modest^78^. They become potentiated by the synergistic combination with MIA. This could be due to the fact that mutated DISC1 might modulate the basal or polyI:C-induced cytokine production by interfering with glycogen synthase kinase-3^79,80^. Alternatively, DISC1 might confer neuronal vulnerability, making them more susceptible to altered microglial function.

The structural (e.g. dendritic arborization, spine density) and functional (e.g. spike rate and timing, oscillatory power) deficits of neonatal dual-hit GE mice were rescued by restoring the normal function of microglia with minocycline – a potent inhibitor of microglial activation and associated neuroinflammation^81^. Minocycline has been found to be neuroprotective in numerous pathologies^82,83^. In particular, its use alone or as adjunctive therapy to antipsychotics improved the behavioral and cognitive performance of schizophrenia patients^84,85^. While the mode of action of minocycline in the adult brain has been well characterized^86^, only few studies focused on its preventive potential during development, before the onset of disease symptoms. When administrated during the course of peripubertal stress exposure, minocycline has been found to prevent the emergence of multiple behavioral abnormalities relevant to human cognitive dysfunction^87^, yet the mechanisms underlying the behavioral rescue are largely unknown. Here, we show that already during neonatal development minocycline is effective in preventing inflammation-induced microglial alteration as well as the associated structural and functional deficits. However, the pathways that selectively link pyramidal neurons in layer II/III of PFC with the anti-inflammatory action of minocycline remain to be investigated in detail. Of note, layer II/III neurons are considerably more dependent of microglia activity than those of layer V/VI^76^.

### Relevance for human mental illness

The relevance of animal models for human mental disorders has often been questioned, since they do not fulfill the validity criteria used for other pathologies. Optimally, animal models recapitulate etiologic processes (i.e. construct validity) or symptom features (i.e. face validity). In case of mental disorders, such as schizophrenia, bipolar disorder or depression, the available mouse models have either excellent construct validity (e.g. models mimicking the genetic background) but limited face validity, or vice versa (e.g models of hippocampal damage or pharmacological models). The dual-hit model used in the present study mimics both genetic and environmental risk factors and recapitulates some of the structural and circuit deficits observed in patients. For example, lower spine density in upper layers of PFC as well as dysfunctional prefrontal gamma band oscillations have been described for schizophrenia patients^17^. Similarly, microglia abnormalities and resulting synaptic deficits have been related to several brain pathologies^76,88^. Therefore, it is tempting to speculate that dual-hit GE mice recapitulate both the etiology (construct validity) as well as the general rules of neuronal, glial, and circuit dysfunction (face validity) that relate to cognitive impairment in mental disorders. They appear highly instrumental for the identification of cellular key players of disease that for ethical and technical reasons are not accessible in humans of comparable age. This brings us closer to one of the major goals of circuit psychiatry that is the identification of key neurobiological targets amenable to tailored therapies^89^ that not only treat but also prevent disease-related cognitive and behavioral deficits.

## METHODS

### Mice

Experiments were performed in compliance with German laws and the guidelines of the European Community for the use of animals in research, and were approved by the local ethical committee (111/12, 132/12). Experiments were carried out on C57Bl/6J mice of both sexes, at the age of P8-10. Heterozygous mutant DISC1 pups carrying a Disc1 allele (Disc1Tm1Kara) on a C57Bl6/J background and C57Bl6/J, whose dams were injected i.v. at embryonic day (E) 9 with the viral mimetic poly I:C (5 mg/kg) were used as dual-hit genetic-environmental model (dual-hit GE)^20^. Genotypes were assessed using genomic DNA (tail biopsies) and following primer sequences: forward primer 5’-TAGCCACTCTCATTGTCAGC-3’ and reverse primer 5’-CCTCATCCCTTCCACTCAGC-3’. Non-treated wild-type mice and the offspring of dams injected at E9 with saline (0.9 %) were used as controls. Timed-pregnant mice from the animal facility of the University Medical Center Hamburg-Eppendorf, of both aforementioned conditions, were housed individually at a 12 h light/12 h dark cycle, and were given access to water and food *ad libitum*. The day of vaginal plug detection was considered as E0.5, while the day of birth as P0.

### Minocycline administration

Minocycline was administered to neonatal mice from P1 to P8 by adding it to the drinking water of the dam, which then passed it on to the offspring via lactation^90^. In line with previous studies^90,91^, the daily dosage of minocycline was 30 mg/kg body weight. To cover the taste of the antibiotic, 8.25 g of sucrose were added to the solution. No difference in liquid intake was observed between dams receiving water and dams receiving water supplemented with minocycline. This administration route has been shown to result in detectable concentrations of the drug in the breast milk of the lactating dam^92^, and to be effective in preventing the adverse effects of MIA in the offspring^93^.

### *In utero* electroporation

Additional wet food supplemented with 2-4 drops of Metacam (meloxicam; 0.5mgml-1; Boehringer-Ingelheim, Germany) was administered from one day before until two days after surgery. At E12.5 or E15.5 randomly assigned pregnant mice received a subcutaneous injection of buprenorphine (0.05 mg/kg body weight) at least 30 min before surgery. Surgery was performed on a heated surface; pain reflexes (toe and tail pinch) and breathing were monitored throughout. Under isoflurane anesthesia (induction: 5%; maintenance: 3.5%), the eyes of the dam were covered with eye ointment to prevent damage, before the uterine horns were exposed and moistened with warm sterile PBS (37 °C). Solution containing 1.25 μg/μl DNA (pAAV-CAG-ChR2(E123T/T159C)-2AtDimer2 or pAAV-CAG-tDimer2)) and 0.1% fast green dye at a volume of 0.75–1.25 μl were injected into the right lateral ventricle of individual embryos using pulled borosilicate glass capillaries with a sharp, long tip. Plasmid DNA was purified with NucleoBond (Macherey-Nagel, Germany). 2A encodes for a ribosomal skip sentence, splitting the fluorescent protein tDimer2 from the opsin during gene translation. Each embryo within the uterus was placed between the electroporation tweezer-type paddles (3 mm diameter for E12.5; 5 mm diameter for E14.5-15.5; Protech, TX, USA) that were roughly oriented at a 20° leftward angle from the midline and a 10° angle downward from anterior to posterior. By these means, neural precursor cells from the subventricular zone, which radially migrate into the medial PFC, were transfected. Electrode pulses (35 V, 50 ms) were applied five times at intervals of 950 ms controlled by an electroporator (CU21EX; BEX, Japan). Uterine horns were placed back into the abdominal cavity after electroporation, which was filled with warm sterile PBS (37 °C). Abdominal muscles and skin were sutured individually with absorbable and non-absorbable suture thread, respectively. After recovery, pregnant mice were returned to their home cages, which were half placed on a heating blanket for two days after surgery. Opsin expression was assessed with a portable fluorescent flashlight (Nightsea, MA, USA) through the intact skull and skin at P2–3 and confirmed post mortem by fluorescence microscopy in brain slices. Pups without expression in the PFC were excluded from the analysis.

### Developmental milestones

Mouse pups were tested for their somatic development and reflexes at P2, P5 and P8. Weight, body and tail length were assessed. Surface righting reflex was quantified as time (max 30 s) until the pup turned over with all four feet on the ground after being placed on its back. Cliff aversion reflex was quantified as time (max 30 s) until the pup withdrew after snout and forepaws were positioned over an elevated edge. Vibrissa placing was rated positive if the pup turned its head after gently touching the whiskers with a toothpick.

### *In vitro* electrophysiology and optogenetics

As previously described^31^, whole-cell patch-clamp recordings were performed from t-Dimer expressing layer II/III and layer V/VI prelimbic neurons in brain slices of P8-10 mice after IUE at E15.5 and E12.5, respectively. Briefly, pups were decapitated, brains were removed and immediately sectioned coronally at 300 **u**m in ice-cold oxygenated high sucrose-based artificial cerebral spinal fluid (ACSF) (in mM: 228 sucrose, 2.5 KCl, 1 NaH2PO4, 26.2 NaHCO3, 11 glucose, 7 MgSO4; 320 mOsm). Slices were incubated in oxygenated ACSF (in mM: 119 NaCl, 2.5 KCl, 1 NaH2PO4, 26.2 NaHCO3, 11 glucose, 1.3 MgSO4; 320mOsm) at 37 °C for 45 min before cooling to room temperature and superfused with oxygenated ACSF in the recording chamber. tDimer2-positive neurons were patched under optical control using pulled borosilicate glass capillaries (tip resistance of 4-7 MΩ) filled with pipette solution (in mM: 130 K-gluconate, 10 HEPES, 0.5 EGTA, 4 Mg-ATP, 0.3 Na-GTP, 8 NaCl; 285mOsm, pH 7.4). Recordings were controlled with the Ephus software in the Matlab environment (MathWorks, MA, USA). Capacitance artefacts and series resistance were minimized using the built-in circuitry of the patch-clamp amplifier (Axopatch 200B; Molecular devices, CA, USA). Responses of neurons to hyper- and depolarizing current injections, as well as blue light pulses (473 nm, 5.2 mW/mm^2^) were digitized at 5 kHz in current-clamp mode.

### *In vivo* electrophysiology and optogenetics

#### Surgery

Multisite extracellular recordings were performed in the PL of P8-10 mice. For recordings in non-anesthetized state, 0.5% bupivacain / 1% lidocaine was locally applied on the neck muscles. For recordings in anesthetized state, mice were injected i.p. with urethane (1 mg/g body weight; Sigma-Aldrich) before surgery. For both groups, the surgery was performed under isoflurane anesthesia (induction: 5%; maintenance: 1.5%). The head of the pup was fixed into a stereotaxic apparatus using two plastic bars mounted on the nasal and occipital bones with dental cement. The bone above the PFC (0.5 mm anterior to bregma, 0.1 mm right to the midline for layer II/III, 0.5 mm for layer V/VI) was carefully removed by drilling a hole of <0.5 mm in diameter. Before recordings, mice were allowed to recover for 10-20 min on a heating blanket.

One- or four-shank multisite optoelectrodes (NeuroNexus, MI, USA) were inserted 2.4 or 1.9 mm (respectively) deep into PFC, perpendicular to the skull surface. One-shank optoelectrodes contained 1×16 recordings sites (0.4-0.8 MΩ impedance, 100μm spacing) aligned with an optical fiber (105 μm diameter) ending 200 μm above the top recording site. Four-shank optoelectrodes contained 4×4 recording sites (0.4-0.8 MΩ impedance, 100 μm spacing, 125 μm intershank spacing) aligned with optical fibers (50 μm diameter) ending 200 μm above the top recording sites. A silver wire was inserted into the cerebellum and served as ground and reference electrode. Before signal acquisition, a recovery period of 15 minutes after electrode insertion was provided.

#### Signal acquisition

Extracellular signals were band-pass filtered (0.1–9,000 Hz) and digitized (32 kHz) with a multichannel extracellular amplifier (Digital Lynx SX; Neuralynx, Bozeman, MO, USA) and the Cheetah acquisition software (Neuralynx). Spontaneous (i.e. not induced by light stimulation) activity was recorded for 15 min at the beginning of each recording session.

#### Light stimulation

Ramp (i.e. linearly increasing power) light stimulations were performed with an arduino uno (Arduino, Italy) controlled diode laser (473 nm; Omicron, Austria). Laser power was adjusted to trigger neuronal spiking in response to >25% of 3-ms-long light pulses at 16 Hz. Resulting light power was in the range of 20–40 mW/mm^2^ at the fiber tip.

#### Post mortem assessment of electrode position

Wide field fluorescence images were acquired to reconstruct the recording electrode position in brain slices of electrophysiologically investigated pups and to localize tDimer2 expression in pups after IUE. Only pups with correct electrode and transfection position were considered for further analysis.

### Histology

#### Perfusion

P8–10 mice were anesthetized with 10% ketamine (aniMedica, Germany) / 2% xylazine (WDT, Germany) in 0.9% NaCl solution (10 μg/g body weight, intraperitoneally (i.p.)) and transcardially perfused with Histofix (Carl Roth, Germany) containing 4% paraformaldehyde.

#### Immunohistochemistry

Brains were postfixed in 4% paraformaldehyde for 24 h and sectioned coronally at 50 μm (immunohistochemistry) or 100 μm (Sholl and spine analysis). Free-floating slices were permeabilized and blocked with PBS containing 0.8% Triton X-100 (Sigma-Aldrich, MO, USA), 5% normal bovine serum (Jackson Immuno Research, PA, USA) and 0.05% sodium azide. Subsequently, slices were incubated overnight with mouse monoclonal Alexa Fluor-488-conjugated antibody against NeuN (1:100, MAB377X; Merck Millipore, MA, USA), rabbit polyclonal primary antibody against CaMKII (1:200, PA5-38239; Thermo Fisher Scientific, MA, USA), rabbit polyclonal primary antibody against GABA (1:1,000, no. A2052; Sigma-Aldrich), or rabbit monoclonal primary antibody against IBA-1 (1:500, catalog #019-19741, Wako), followed by 2 h incubation with Alexa Fluor-488 goat anti-rabbit IgG secondary antibody (1:500, A11008; Merck Millipore). Finally, slices were transferred to glass slides and covered with Vecta-Shield (Vector Laboratories).

#### Imaging

Sections were examined with a confocal microscope (DM IRBE, Leica Microsystems and Zeiss LSN700). To quantify the t-Dimer overlap with NeuN, CaMKII and GABA, microscopic fields over PFC were acquired as 1024×1024 pixel images (pixel size, 1465 nm) using a 10X objective (numerical aperture, 0.3). The same settings were used to quantify the number of CaMKII positive neurons (n=4 fields per section, 3 sections per mouse). For IBA-1, 20-images microscopic stacks (n=8 stacks per section, 3 sections per mouse) were acquired as 512×512 pixel images (pixel size, 732 nm; Z-step, 1000 nm) using a 40X objective (numerical aperture, 1.25). Microscopic stacks used for Sholl and spine analysis were acquired as 2048×2048 pixel images (pixel size, 156 nm; Z-step, 1000 and 500 nm, respectively).

### Image analysis

#### CaMKII+ cells quantification

The number of CaMKII-positive neurons was semi-automatically assessed with a custom-written algorithm in the ImageJ environment. Briefly, a Region of Interest (ROI) was manually placed over either superficial or deep prefrontal layers. The image contrast was enhanced (*enhance contrast* function, 0.5% of saturated pixels) and a *median filter* was applied (radius=1.5). To reduce background noise, we used the *subtract background* function, with a radius of 30 pixels. The image was then binarized (*convert to mask*) and segmented using the *watershed* function. To identify the neurons we used the *extended maxima* function of the MorphoLibJ^94^ plugin (dynamic=30, connectivity=4). We subtracted the regional maxima with the lowest intensity (i.e. the objects with bigger area) using *area opening* (pixel=150) and counted the remaining objects (*analyze particles*).

#### Neuronal morphological analysis

Sholl analysis and spine density quantification were carried out in the ImageJ environment. For Sholl analysis, images were binarized (*auto threshold*) and dendrites were traced using the semi-automatical plugin *Simple Neurite Tracer*. The traced dendritic tree was analyzed with the plugin *Sholl Analysis*, after the geometric center was identified using the *blow/lasso tool*. For spine density quantification, we first traced the dendrite of interest (apical, basal, proximal oblique or secondary apical) and measured its length (*line*) and then manually counted the number of spines (*point picker*).

#### IBA-1+ cells quantification

To quantify the number of IBA-1 stained cells we used a custom-written algorithm in ImageJ. The image stacks were collapsed to a maximum intensity Z-projection, and background noise was subtracted (*despeckle*). To facilitate automatic thresholding, the image was passed through a gaussian filter (*gaussian blur*, sigma=2) before being binarized (*auto threshold* with the *triangle* method). The number of cells was counted using *analyze particles* (size>150 pixels).

#### IBA-1+ cells morphological analysis

The morphology of microglial cells was assessed on maximum intensity Z-projections in the Matlab environment, using previously reported criteria^95,96^. Images were automatically thresholded (*graythresh* and *im2bw* functions) and putative microglial cells were identified as objects between 200 and 1500 pixels (*bwareaopen*). Around the center of mass of each of the isolated cells, a region of interest (ROI) of 110×110 pixels was computed and visually examined. If the ROI contained a properly segmented microglia cells, its features (area, perimeter, eccentricity) were quantified (*regionprops*). ROIs in which the microglial cell touched the boundaries of the image or in which more than one cell was included were discarded. Further, cellspread (analogous to process length) was computed as the average distance between the center of mass and the “extrema” of the cell; roundness was defined as the ratio between 4*pi*area and the square of the perimeter of the cell.

### *In vitro* electrophysiology

As previously described^31^, data were imported and analyzed offline using custom-written tools in the Matlab environment (MathWorks). For *in vitro* data, all potentials were corrected for liquid junction potentials (−10 mV) for the gluconate-based electrode solution. The RMP was measured immediately after obtaining the whole-cell configuration. To assess input resistance, hyperpolarizing current pulses of 200 ms duration were applied. Active membrane properties and current–voltage relationships were determined by unsupervised analysis of responses to a series of 600 ms long hyper- and depolarizing current pulses. Amplitude of APs was measured from threshold to peak.

### *In vivo* electrophysiology

*In vivo* data were analyzed with custom-written algorithms in the Matlab environment. Data were processed as following: band-pass filtered (500–5,000 Hz) to analyze MUA and bandpass filtered (4-100 Hz) using a third-order Butterworth filter before downsampling to 3.2 kHz to analyze LFP. All filtering procedures were performed in a phase preserving manner.

#### Detection of oscillatory activity

The detection of discontinuous patterns of activity in the neonatal PL were performed using a modified version of the previously developed algorithm for unsupervised analysis of neonatal oscillations^26^. Briefly, deflections of the root mean square of band-pass filtered signals (1–100 Hz) exceeding a variance-depending threshold were considered as network oscillations. The threshold was determined by a Gaussian fit to the values ranging from 0 to the global maximum of the root-mean-square histogram. If two oscillations occurred within 200 ms of each other they were considered as one. Only oscillations lasting > 1 s were included, and their occurrence, duration and amplitude were computed.

#### Power spectral density

For power spectral density analysis, 1 s-long windows of network oscillations were concatenated and the power was calculated using Welch’s method with non-overlapping windows. For optical stimulation, we compared the average power during the 1.5 s-long time window preceding the stimulation to either the first or to the last 1.5 s-long time window of light-evoked activity.

#### Multi-unit activity

MUA was detected as the peak of negative deflections exceeding five times the standard deviation of the filtered signal and having a prominence larger than half the peak itself.

#### Single unit activity

SUA was detected and clustered using Offline Sorter (Plexon, TC, USA). 1–4 single units were detected at each recording site. Data were imported and analyzed using custom-written tools in the Matlab software (MathWorks).

#### Firing rate

The firing rate was computed by dividing the total number of spikes by the duration of the analyzed time window.

#### Inter-spike-interval

Inter-spike interval (ISI) was calculated at 1 ms resolution.

#### Phase-locking

To quantify locking of spiking activity to different cortical rhythms, we first bandpass-filtered the LFP in the frequency band of interest, and extracted the phase by using the Hilbert transform. Subsequently, the phase-locking value (PLV) was computed (*circ_r*).

#### Pairwise phase consistency

Pairwise phase consistency (PPC) was computed as previously described^97^. Briefly, the phase in the band of interest was extracted as mentioned above, and the mean of the cosine of the absolute angular distance (dot product) among all pairs of phases was calculated.

#### Spike-triggered LFP power

Spike-triggered LFP spectra were calculated as

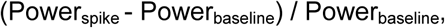

where the spike-triggered power spectrum (Power_spike_) was calculated using Welch’s method for a 200 ms-log time window centered on each spike, and the power spectrum of baseline LFP (Power_baseline_) was averaged for two time windows, 100-300 ms and 200-400 ms before each spike.

## Statistical analysis

Statistical analyses were performed using R Statistical Software (Foundation for Statistical Computing, Vienna, Austria). Data were tested for normality with the Shapiro-Wilk test, for homoscedasticity with the Breush-Pagan test, and for equal variances with the Bartlett’s test. Normally distributed, homoscedastic, having equal variance and non-nested data were tested for significant differences (*P<0.05, **P<0.01 and ***P<0.001) using paired t-test, unpaired t-test or one-way repeated-measures analysis of variance with Bonferroni-corrected post hoc analysis. Not normally distributed, heteroskedastic or not having equal variance data were tested with yuen’s bootstrap test with 5000 repetitions, robust one-way repeated measures ANOVA with Bonferroni-Holm corrected post hoc analysis or robust repeated measures ANCOVA with age as covariate and Bonferroni-Holm corrected post hoc analysis (*yuenbt*, *rmanova* and *ancova* functions of the WRS2-R package^98^). A standard 20% level of trimming for the mean was selected for these tests. Such tests were preferred to more traditional non-parametric tests in virtue of the (sometimes) high levels of unequal variance in our data^99^. To account for the commonly ignored increased false positive rate inherent in nested design^100^, nested data were analyzed with linear mixed-effect models. Parameter estimation was done using the *lmer* function implemented in the lme4-R package^101^. Model selection was performed using the Akaike Information Criterion (AIC) and/or the Bayesian information criterion (BIC), as differences between the two criteria were minimal. To test the significance of condition in our model, we performed a likelihood ratio test against a reduced model in which we removed condition. The circular statistics toolbox was used to test for significant differences in the phase locking data. No statistical measures were used to estimate sample size since effect size was unknown. Investigators were blinded to the group allocation when Sholl and spine analyses were performed. Unsupervised analysis software was used to preclude investigator biases. Statistical parameters can be found in the main text and/or in the figure legends.

## Data and code availability

The authors declare that all data and code supporting the findings of this study are included in the manuscript and its Supplementary Information or are available from the corresponding author on request.

## Acknowledgments

We thank Dr. Joseph Gogos for providing the DISC1 mice, Drs. S. Wiegert and T. Oertner for providing opsin derivates, and A. Marquardt, A. Dahlmann, I. Ohmert, and K. Titze for excellent technical assistance. This work was funded by grants from the European Research Council (ERC-2015-CoG 681577 to I.L.H.-O.) and the German Research Foundation (Ha 4466/10-1, SPP 1665, SFB 936 B5 to I.L.H.-O. and C6 to C.M.).

I.L. H.-O. is member of FENS Kavli Network of Excellence.

